# Applying generalised allometric regressions to predict live body mass of tropical and temperate arthropods

**DOI:** 10.1101/297697

**Authors:** Esra H. Sohlström, Lucas Marian, Andrew D. Barnes, Noor F. Haneda, Stefan Scheu, Björn C. Rall, Ulrich Brose, Malte Jochum

## Abstract

1. The ecological implications of body size extend from the biology of individual organisms to ecosystem–level processes. Measuring body mass for high numbers of invertebrates can be logistically challenging, making length-mass regressions useful for predicting body mass with minimal effort. However, standardised sets of scaling relationships covering a large range in body length, taxonomic groups, and multiple geographical regions are scarce.

2. We collected 6293 arthropods from 19 higher-level taxa in both temperate and tropical locations to compile a comprehensive set of linear models relating live body mass to a range of predictor variables. For each individual, we measured live weight (hereafter, body mass), body length and width, and conducted linear regressions to predict body mass using body length, body width, taxonomic group and geographic region. Additionally, we quantified prediction error when using parameters from arthropods of a different geographic region.

3. Incorporating body width into taxon- and region-specific length-mass regressions yielded the highest prediction accuracy for body mass. Using regression parameters from a different geographic location increased prediction error, causing over- or underestimation of body mass depending on geographical origin and whether body width was included.

4. We present a comprehensive range of parameters for predicting arthropod body mass and provide guidance for selecting optimal scaling relationships. Given the importance of body mass for functional invertebrate ecology and a paucity of adequate regressions to predict arthropod body mass from different geographical regions, our study provides a long-needed resource for quantifying live body mass in invertebrate ecology research.

## Introduction

Body size is one of the most fundamental traits of living organisms (Peters, 1983). From the individual to the community level, a vast range of ecosystem properties scale with arthropod body size. Body size determines various aspects of an organism’s individual biology, such as life-history, behaviour, range size, movement and physiology (Bekoff *et al.*, 1981; Woodward *et al.*, 2005; White *et al.*, 2007; Hirt *et al.*, 2017). Aspects shaping arthropod communities such as species abundance, biomass production, trophic link structure, and interaction strengths are also related to the body size of constituent individuals and populations (Boudreau *et al.*, 1991; Belgrano *et al.*, 2002; Brose *et al.*, 2006; Riede *et al.*, 2011; Rall *et al.*, 2012; Kalinkat *et al.*, 2013). As a result, arthropod body size determines how individuals and communities carry out functions, making it a powerful predictor of ecosystem performance (Barnes *et al.*, 2018).

Most biological rates scale with body size following a power-law relationship (Peters, 1983; White *et al.*, 2007), which has important implications for individual and community ecology. In the early 1930s, Kleiber (1932) proposed an allometric scaling relationship of metabolism with body mass following a ¾ power law function, though this has been extensively debated (see Brown *et al.*, 2004; Kolokotrones *et al.*, 2010; Ehnes *et al.*, 2011).This power-law scaling means that smaller animals have a lower per capita metabolic rate than larger ones, though their mass-specific metabolic rate is higher, yielding distinct patterns of energy demand in populations and communities depending on the relationship between body size and total biomass (Reichle, 1968). Additionally, home- and foraging ranges of animals increase with body size, which has been demonstrated for a wide range of organisms, from small invertebrates to large mammals (Lindstedt *et al.*, 1986; Swihart *et al.*, 1988; Jetz *et al.*, 2004; Greenleaf *et al.*, 2007). Due to the allometric scaling of a large range of physiological and ecological properties, one can utilise general scaling relationships to predict ecological properties from measured values of organism body size.

While body size is highly useful as a predictive trait for both the response of arthropods to environmental change and also their effects on ecosystem processes, ecologists face many logistic challenges when collecting body size data. Firstly, measurement of arthropod body mass is particularly challenging due to their small body size and typically high abundance. As a consequence, researchers might measure a few individuals of each species and apply an average of these values to the remaining individuals. This practice eliminates intraspecific variation that occurs among sampling sites, especially when the sites are distributed along ecological gradients (Violle *et al.*, 2012). However, in large field sampling campaigns, collecting individual body mass data across all samples is often infeasible due to the logistic difficulties of weighing large numbers of individual organisms. Furthermore, data on live—rather than dry—body mass is often required to accurately relate body size to a range of ecological attributes. For example, physiological rates (such as metabolism), species interactions (e.g., pollination and predation), and behavioural patterns are directly dependent on the body mass of an animal while it is alive, as opposed to its dry mass which serves only as a proxy for live mass. However, dry mass is far more frequently measured as it is extremely difficult to take live body mass measurements of arthropods, particularly in large sampling campaigns. This limitation calls for the provision of practical and accurate tools to acquire individual-level, live arthropod body mass data in order to assess population and community responses in arthropod size structure and investigate corresponding ecosystem processes.

Length-mass regressions have proven to be a powerful tool to predict body mass based on body length measurements (Rogers *et al.*, 1977; Schoener, 1980; Benke *et al.*, 1999; Johnston & Cunjak, 1999; Gruner, 2003; Wardhaugh, 2013) which are much easier to obtain than measurements of body mass. This approach relies on regression parameters estimated for length-mass relationships, which can be used to predict body mass when only body length data are available. However, finding suitable regression parameters for a given dataset (for example, where parameters are from the same taxonomic group and geographical region) is often not possible. This limitation can be problematic because scaling relationships—and thus, their regression parameters—are likely to vary substantially among taxonomic groups and geographic regions; a discrepancy that has been shown to be especially distinct between tropical and temperate regions (Schoener, 1980; Gruner, 2003; Wardhaugh, 2013). Thus, using length–mass regression parameters from a different geographical region is likely to increase the error in predictions of body mass. Finally, to our knowledge there are no regressions available in the literature that are based on live body-mass measurements and that cover a large range of taxa and multiple geographic regions. Therefore, researchers are typically constrained to using rough conversion factors (Peters, 1983) or more elaborate dry mass–fresh mass regressions (e.g. Mercer *et al.*, 2001), which add further error to body mass predictions due to the very same sources of variation in length-mass scaling relationships (geographic origin, taxon-specificity, etc.). Considering the broad application of body-size data in ecological research, there are surprisingly few studies that provide length-mass regression parameters for terrestrial arthropods, and these are generally restricted to one of either temperate or tropical animals, or to only a few taxonomic groups (Schoener, 1980; Burgherr & Meyer, 1997; Benke *et al.*, 1999; Gruner, 2003; Wardhaugh, 2013).

In this paper, we provide an unprecedented dataset of length–mass scaling relationships based on measurements of live body mass and body length of 6293 terrestrial arthropods from both tropical and temperate geographical regions. We measured body mass while the animals were still alive. As such, our regressions will be particularly useful for researchers interested in the physiology, behavior or interaction ecology of their target organisms. We performed length–mass regressions for arthropods, including various combinations of body width, taxonomic group and geographic origin as additional co-variables, and compared the accuracy in predicting body mass among these various models. We hypothesised that prediction accuracy improves with an increasing number of additional predictors (e.g., including body width, taxonomic group and geographic region), as opposed to using only body length as a sole predictor of body mass. Additionally, we expected a higher prediction accuracy when using regression parameters taken from the same geographic region, as opposed to using regression parameters of arthropods from a different geographic region (hereafter, geographically-disjunct regression parameters). Our study thus provides a generalised resource for predicting live body mass across an unprecedented range of terrestrial arthropod groups (including 19 orders of Arachnida, Myriapoda, Crustacea and Insecta), as well as guidance for deciding which scaling relationships to use for predicting arthropod body mass depending on the dataset at hand.

## Materials and Methods

### Study sites and sampling techniques

To account for different scaling relationships in temperate versus tropical geographical regions, we chose two sampling locations: one temperate location in Germany and one tropical location in Indonesia. Temperate sites were located near Göttingen, Germany (51°32′02″N, 09°56′08″E) at an altitude of around 150 m asl, with a mean annual air temperature of 7.4 °C, mean annual precipitation of 700 mm (Heinrichs *et al.*, 2014) and a vegetation growth period from May to September. Tropical sites were located near Jambi City in Sumatra, Indonesia (1°35′24″S 103°36′36″E), at an altitude around 20 m asl. Jambi City has a mean annual air temperature of 25 °C and a mean annual precipitation of 2100 to 2800 mm (Ishizuka *et al.*, 2002). The sampling sites in both regions included wayside vegetation, open grassland areas and forest strips. Sampling sites were chosen due to their proximity to the laboratory in both regions to ensure a fast and simple work flow, since animals had to be kept alive after collection and living animals could not be stored for more than eight hours to avoid increased body mass-loss.

Three standard sampling techniques were used in order to cover a broad variety of arthropod taxa and to achieve a sufficient overlap of taxonomic groups from both sampling regions. For active and fast moving ground animals, as well as nocturnal species, live pitfall traps (diameter of 11 cm and height of 12 cm) were used within forest and grassland sites. Pitfall traps were closed with a funnel-shaped lid to prevent animals from escaping. Pitfall traps were buried so the opening of the pitfall was flush with the surface of the ground. They were installed in the morning and animals were collected after 24 hours to avoid loss of individuals due to predation, drowning, or desiccation. Sweep nets were used in open grassland and wayside vegetation plots to collect animals from within low vegetation, shrubs and small trees to sample stationary, as well as fast-moving and flying animals. At the forest sites, less mobile animals from within the litter layer were collected via leaf-litter sieving. Material from the loose leaf litter (F-Layer) on top of the humus layer was collected and sieved with a coarse-meshed grid (2 × 2 cm). Animals that fell through the mesh were hand-collected from a collecting tray and stored in individual vials for further processing.

### Morphological measurements and data collection

Arthropods were stored in a refrigerator at 10 °C for a maximum of 8 hours after collection to slow down their metabolism and reduce body mass loss. In order to maximise accuracy in live body mass measurements, we conducted preliminary tests of body mass-loss following live capture, comparing live to recently killed arthropods to establish whether specimens should be weighed when alive or dead. As we found considerable variation in body mass between live and dead animals, we weighed all arthropods whilst still alive on a precision scale (to the nearest 0.01 mg) and subsequently stored them in ethanol (75 %). For measurements of length and maximum width (to the nearest 0.01 mm), pictures of the dorsal or ventral and lateral view were taken with a Dino-Lite Digital Microscope (Dino-Lite Edge; AnMo Electronics Corporation). Afterwards, each individual was measured using ImageJ (Version 1.48k or newer), leaving out appendages to generalize the process. Finally, every individual was identified to family level using ‘Insects of Australia’ (Commonwealth Scientific and Industrial Research Organization (Australia), 1991), ‘Spider Families of the World’ (Jocqué & Dippenaar-Schoeman, 2007) and the identification keys of ‘Brohmer – Fauna von Deutschland’(Schaefer, 2009). All underlying data can be found online in the Supporting Information (Supporting Data S1).

### Statistical analysis

All statistical analyses were performed using R Version 3.4.0 (R Core Team, 2015). Prior to the analysis, raw data of body length, mass and width were log_10_-transformed. Taxa without width measurements were excluded from the main analysis. However, length-mass regressions for these taxonomic groups, along with a range of regressions for higher-resolution taxonomic groups, were carried out separately and results are presented in the Supporting Information (i.e., regressions for selected taxa based on morphology, taxonomy or behaviour; Table S1).

We performed linear models to test the relationship between body mass and length (L) alone, and with the co-variables width (W), taxonomic group (T) and geographical region (R) in all possible combinations, yielding eight linear models in total (Table 1). For models that included taxonomic group and geographical region, these two factors were combined and treated as a single factor (e.g. temperate Araneae or tropical Araneae to account for the uneven distribution of some taxonomic groups across geographical regions. The most complex model included length, width, taxonomic group and geographic region (model LWTR) and the least complex model included only length as a single independent variable (model L) (see Supporting Methods S1 for a worked example of body mass predictions using each model type). Finally, model fits were then compared using Akaike’s Information Criterion (AIC).

**Table 1:** Model comparisons for the eight linear models used to predict live body mass based on different explanatory variables. Models are compared based on AIC and r^2^.

| Model no. | Model parameters | AIC | $\Delta$ AIC | $r^2$ |
| --- | --- | --- | --- | --- |
| 1 (LWTR) | Length, width, taxa, region | -8551.2011 | 0 | 0.9860 |
| 2 (LWT) | Length, width, taxa | -8087.7384 | 463.4627 | 0.9849 |
| 3 (LWR) | Length, width, region | -4377.9316 | 4173.2695 | 0.9725 |
| 4 (LW) | Length, width | -4267.3045 | 4283.8966 | 0.9438 |
| 5 (LTR) | Length, taxa, region | -1179.7326 | 7371.4685 | 0.9546 |
| 6 (LT) | Length, taxa | -793.3381 | 7757.8630 | 0.9516 |
| 7 (LR) | Length, region | 3050.0298 | 11601.2309 | 0.9103 |
| 8 (L) | Length | 3249.5840 | 11800.7851 | 0.8143 |

Because we hypothesised that using regression parameters from different geographic regions likely increases error in predictions of arthropod body mass, we assessed this prediction error by quantifying the proportional difference between predicted and observed body mass using geographically-disjunct and geographically non-disjunct regression parameters for the two all-taxa models (models LWR and LR). Specifically, body mass prediction accuracy of regression parameters was calculated as the log-response ratio

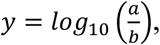

where *y* is the prediction error of body mass, *a* is the predicted body mass using length-mass regressions and *b* is observed body mass. We then assessed how prediction accuracy varied across the range of body length to ascertain if there might be systematic error in body mass predictions depending on arthropod body size.

## Results

In total, 6293 individuals from 19 arthropod higher-order taxa were collected, weighed while alive, and measured for body length and width across the Indonesian and German sites (hereafter, tropical and temperate geographic regions). Body length of collected arthropods ranged from 0.60 mm to 68.12 mm and body mass ranged from 0.01 mg to 5108.57 mg (Table 2). As expected, we found a consistent positive scaling relationship for body mas with body length across all collected arthropods.

**Table 2:** Taxonomic groups sampled in the two geographic regions (temperate and tropical), including the number of individuals (n), number of families, length range and mass range (live body mass) per taxon.

| Order | n |  | No. of families |  | Length range (mm) |  | Mass range (mg) |  |
| --- | --- | --- | --- | --- | --- | --- | --- | --- |
|  | Temp. | Trop. | Temp. | Trop. | Temp. | Trop. | Temp. | Trop. |
| Araneae | 519 | 1081 | 16 | 27 | 1.01 - 12.26 | 0.78 - 25.71 | 0.15 - 212.78 | 0.01 - 5108.57 |
| Coleoptera | 408 | 298 | 15 | 21 | 1.66 - 35.10 | 1.10 - 43.42 | 0.33 - 1067.93 | 0.05 - 3698.96 |
| Dermaptera | 60 | 130 | 2 | 3 | 3.00 - 13.96 | 1.87 - 18.71 | 2.13 - 72.06 | 0.01 - 92.57 |
| Dictyoptera | - | 247 | 1 | 6 | - | 1.69 - 65.07 | - | 0.42 - 1060.93 |
| Diptera | 504 | 189 | 31 | 28 | 1.49 - 16.82 | 1.58 - 23.61 | 0.07 - 74.50 | 0.07 - 165.17 |
| Geophilomorpha | - | 13 | - | 2 | - | 7.47 - 33.54 | - | 0.29 - 21.03 |
| Hemiptera | 598 | 454 | 14 | 35 | 1.31 - 12.05 | 0.95 - 23.76 | 0.27 - 146.90 | 0.05 - 261.53 |
| Hymenoptera | 222 | 371 | 14 | 23 | 1.70 - 22.26 | 0.62 - 31.88 | 0.06 - 835.43 | 0.01 - 1664.61 |
| Isopoda | 88 | 88 | 6 | 3 | 2.45 - 16.16 | 2.45 - 16.16 | 0.81 - 181.27 | 0.22 - 189.52 |
| Lepidoptera | 31 | 121 | 4 | 10 | 3.56 - 16.23 | 3.23 - 35.52 | 1.67 - 91.02 | 0.56 - 908.65 |
| Lithobiomorpha | 161 | 60 | 1 | 1 | 2.77 - 23.63 | 2.22 - 51.21 | 0.65 - 170.65 | 0.01 - 439.53 |
| Neuroptera | 21 | 18 | 2 | 4 | 3.79 - 11.34 | 3.26 - 27.29 | 2.61 - 17.44 | 1.33 - 144.05 |
| Odonata | - | 21 | - | 4 | - | 23.37 - 54.83 | - | 14.24 - 367.32 |
| Opiliones | 89 | 24 | 3 | 3 | 0.93 - 7.53 | 1.09 - 10.09 | 0.81 - 95.02 | 0.40 - 165.61 |
| Orthoptera | 35 | 277 | 2 | 6 | 3.79 - 24.28 | 1.28 - 68.12 | 3.81 - 417.84 | 0.14 - 3895.10 |
| Polydesmida | 12 | 80 | 1 | 1 | 9.21 - 19.95 | 4.02 - 32.55 | 9.24 - 67.25 | 0.05 - 205.02 |
| Pseudoscorpionida |  | 36 | - | 2 | - | 0.95 - 4.16 | 1.33 - 19.91 | 0.16 - 2.12 |
| Psocoptera |  | 26 | - | 3 | - | 1.12 - 2.92 | 0.22 - 0.64 | 0.11 - 8.00 |
| Scolopendromorpha | - | 11 | - | 2 | - | 4.83 - 41.84 | - | 0.88 - 276.18 |
| Total (geogr. region) | 2748 | 3545 | 122 | 190 | 0.930 - 35.1 | 0.62 - 68.12 | 0.06 - 1067.93 | 0.01 - 5108.57 |
| **Grand total** | **6293** |  | **246** |  | **0.60 - 68.10** |  | **0.01 - 5108.57** |  |

The most complex model (Model LWTR, including body length, body width, taxonomic group and geographic region as predictors) best explained variation in body mass according to AIC selection and r^2^ (Table 3). The consistently positive slope in the relationship between body length and body mass (for all arthropod taxa except for Odonata and Neuroptera) was significantly influenced by body width, taxonomic group and geographic region that the arthropods originated from (Table 3, Figure 1). Thus, the slope of the length-mass relationship varied with body width, taxonomic group and geographic region (e.g. the slope of the length-mass relationship differed between spiders and beetles as well as between temperate and tropical spiders).

**Table 3:**
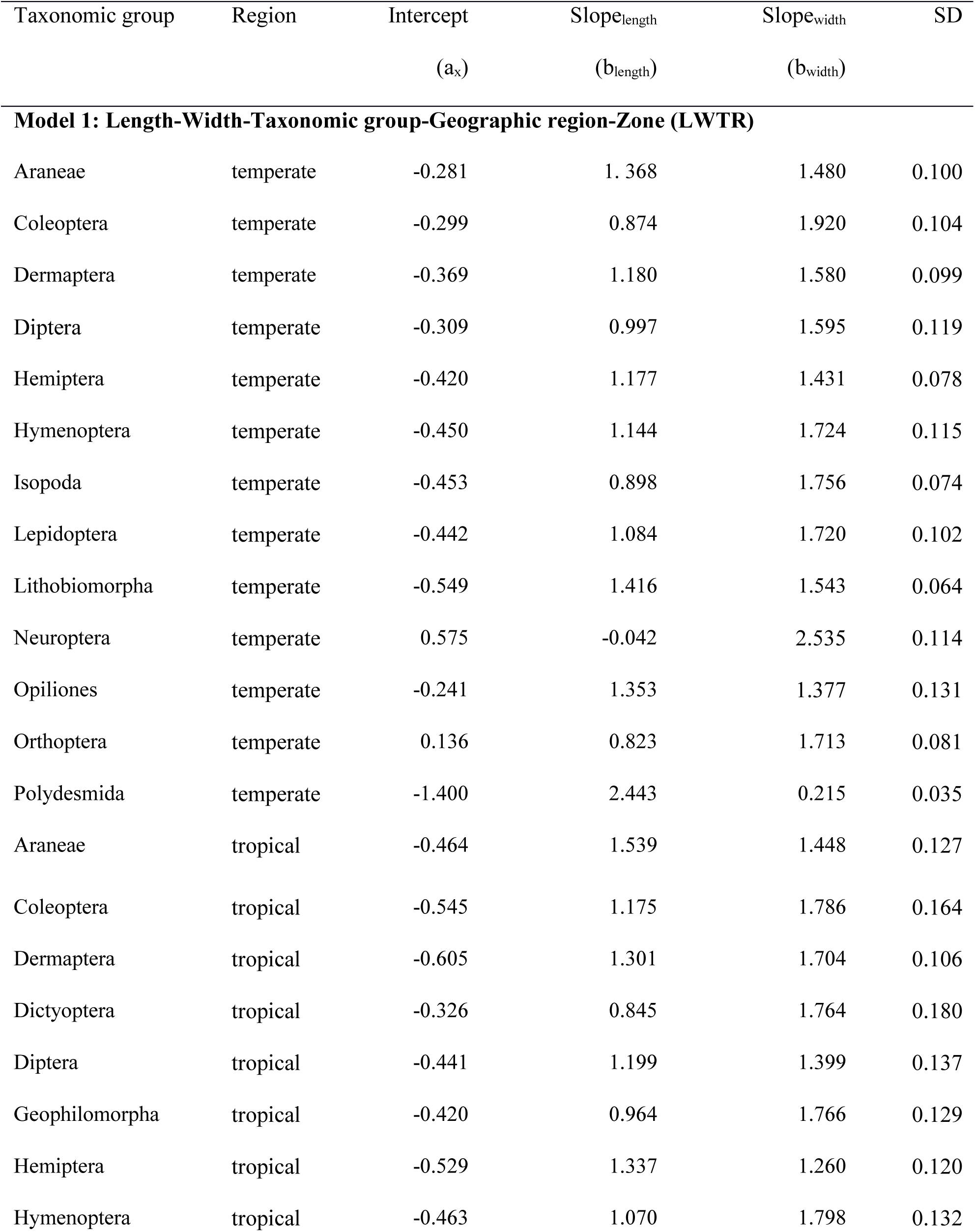
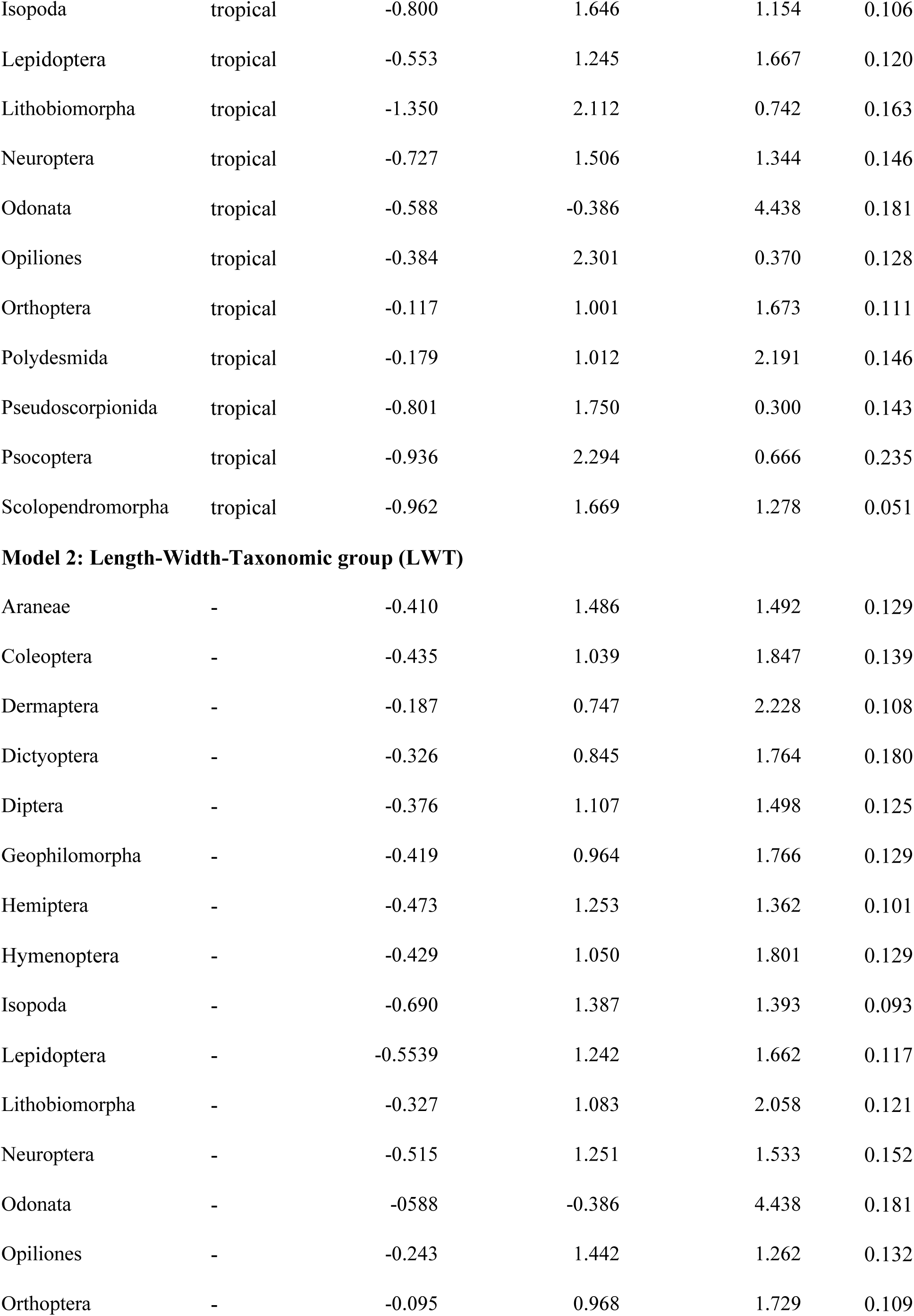
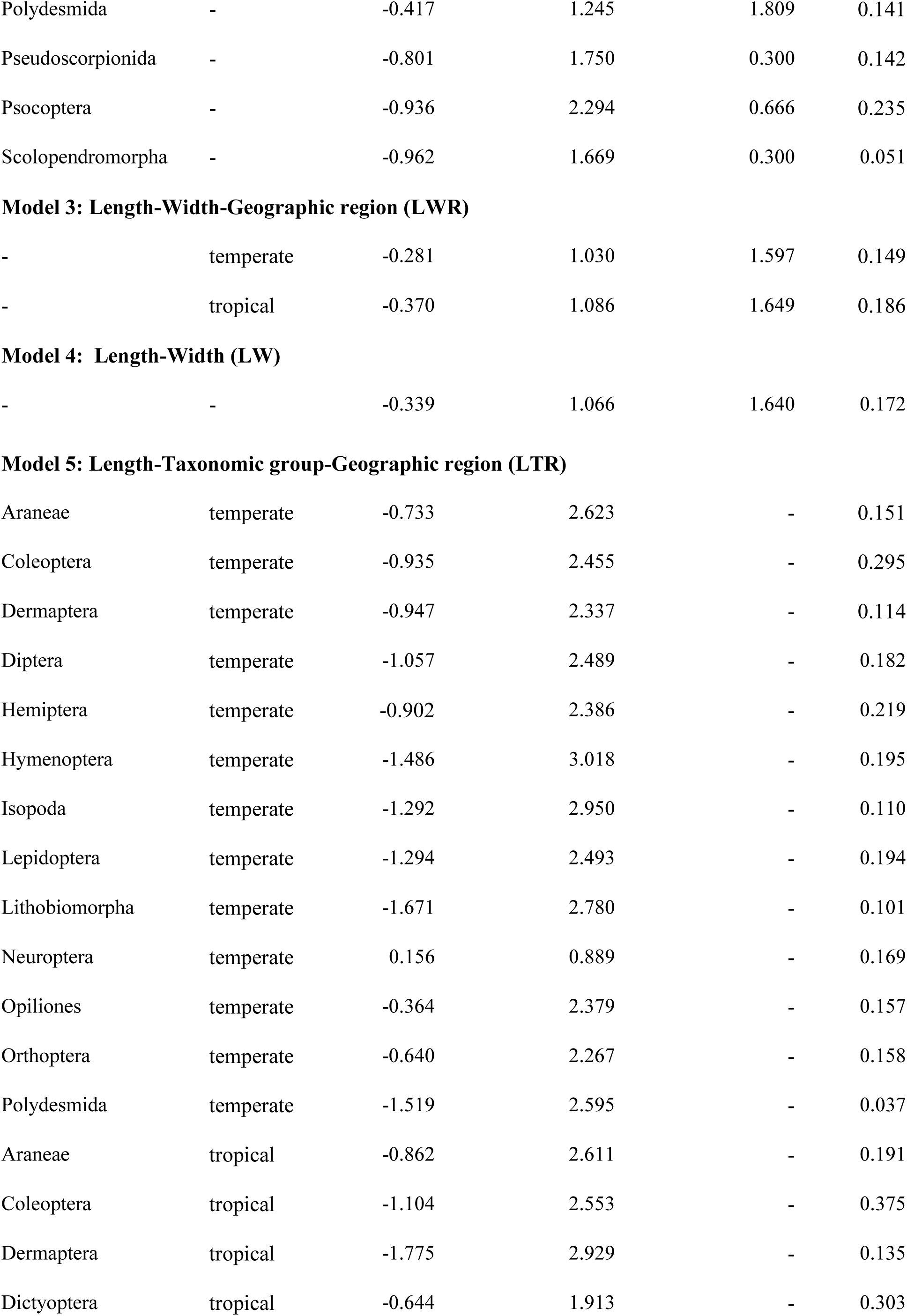
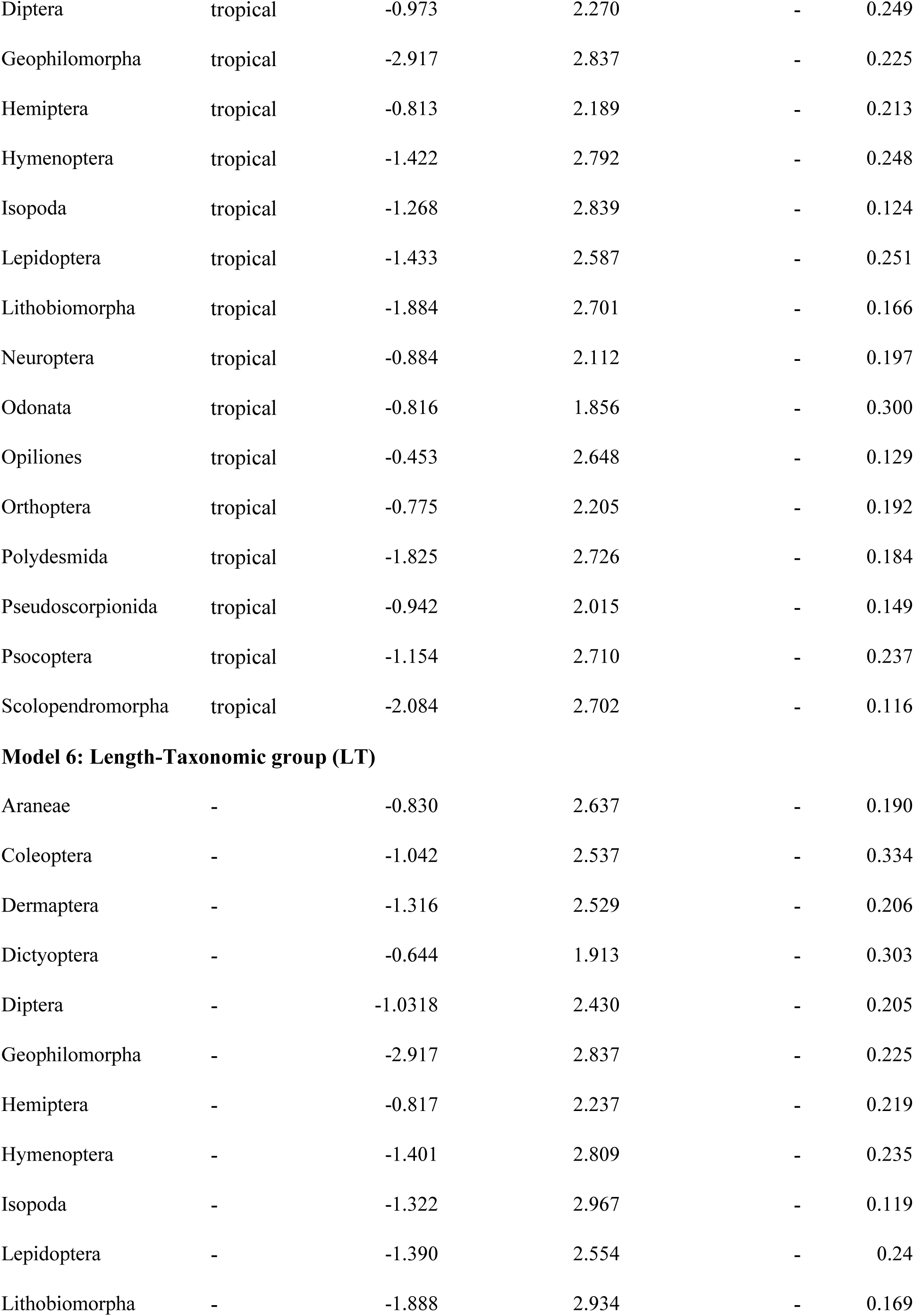
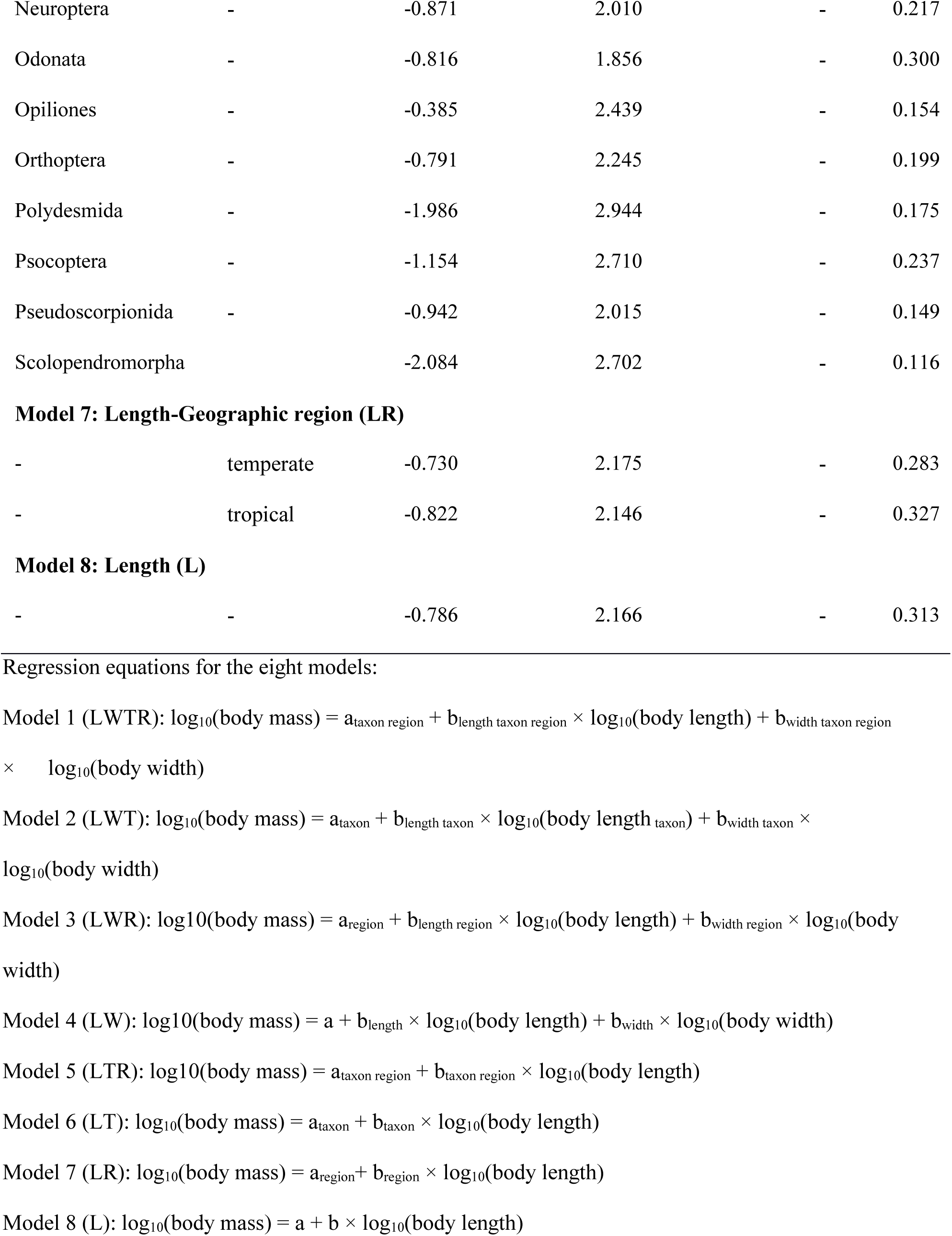
Regression parameters for the eight linear models for live body mass prediction in dependence of body length (L, in mm), maximum body width (W, in mm), taxonomic group (T) and geographic region (R, temperate and tropical).

| Taxonomic group | Region | Intercept<br>($a_x$) | Slope<sub>length</sub><br>($b_{\text{length}}$) | Slope<sub>width</sub><br>($b_{\text{width}}$) | SD |
| --- | --- | --- | --- | --- | --- |
| **Model 1: Length-Width-Taxonomic group-Geographic region-Zone (LWTR)** |  |  |  |  |  |
| Araneae | temperate | -0.281 | 1.368 | 1.480 | 0.100 |
| Coleoptera | temperate | -0.299 | 0.874 | 1.920 | 0.104 |
| Dermaptera | temperate | -0.369 | 1.180 | 1.580 | 0.099 |
| Diptera | temperate | -0.309 | 0.997 | 1.595 | 0.119 |
| Hemiptera | temperate | -0.420 | 1.177 | 1.431 | 0.078 |
| Hymenoptera | temperate | -0.450 | 1.144 | 1.724 | 0.115 |
| Isopoda | temperate | -0.453 | 0.898 | 1.756 | 0.074 |
| Lepidoptera | temperate | -0.442 | 1.084 | 1.720 | 0.102 |
| Lithobiomorpha | temperate | -0.549 | 1.416 | 1.543 | 0.064 |
| Neuroptera | temperate | 0.575 | -0.042 | 2.535 | 0.114 |
| Opiliones | temperate | -0.241 | 1.353 | 1.377 | 0.131 |
| Orthoptera | temperate | 0.136 | 0.823 | 1.713 | 0.081 |
| Polydesmida | temperate | -1.400 | 2.443 | 0.215 | 0.035 |
| Araneae | tropical | -0.464 | 1.539 | 1.448 | 0.127 |
| Coleoptera | tropical | -0.545 | 1.175 | 1.786 | 0.164 |
| Dermaptera | tropical | -0.605 | 1.301 | 1.704 | 0.106 |
| Dictyoptera | tropical | -0.326 | 0.845 | 1.764 | 0.180 |
| Diptera | tropical | -0.441 | 1.199 | 1.399 | 0.137 |
| Geophilomorpha | tropical | -0.420 | 0.964 | 1.766 | 0.129 |
| Hemiptera | tropical | -0.529 | 1.337 | 1.260 | 0.120 |
| Hymenoptera | tropical | -0.463 | 1.070 | 1.798 | 0.132 |
| Isopoda | tropical | -0.800 | 1.646 | 1.154 | 0.106 |
| Lepidoptera | tropical | -0.553 | 1.245 | 1.667 | 0.120 |
| Lithobiomorpha | tropical | -1.350 | 2.112 | 0.742 | 0.163 |
| Neuroptera | tropical | -0.727 | 1.506 | 1.344 | 0.146 |
| Odonata | tropical | -0.588 | -0.386 | 4.438 | 0.181 |
| Opiliones | tropical | -0.384 | 2.301 | 0.370 | 0.128 |
| Orthoptera | tropical | -0.117 | 1.001 | 1.673 | 0.111 |
| Polydesmida | tropical | -0.179 | 1.012 | 2.191 | 0.146 |
| Pseudoscorpionida | tropical | -0.801 | 1.750 | 0.300 | 0.143 |
| Psocoptera | tropical | -0.936 | 2.294 | 0.666 | 0.235 |
| Scolopendromorpha | tropical | -0.962 | 1.669 | 1.278 | 0.051 |
| **Model 2: Length-Width-Taxonomic group (LWT)** |  |  |  |  |  |
| Araneae | - | -0.410 | 1.486 | 1.492 | 0.129 |
| Coleoptera | - | -0.435 | 1.039 | 1.847 | 0.139 |
| Dermaptera | - | -0.187 | 0.747 | 2.228 | 0.108 |
| Dictyoptera | - | -0.326 | 0.845 | 1.764 | 0.180 |
| Diptera | - | -0.376 | 1.107 | 1.498 | 0.125 |
| Geophilomorpha | - | -0.419 | 0.964 | 1.766 | 0.129 |
| Hemiptera | - | -0.473 | 1.253 | 1.362 | 0.101 |
| Hymenoptera | - | -0.429 | 1.050 | 1.801 | 0.129 |
| Isopoda | - | -0.690 | 1.387 | 1.393 | 0.093 |
| Lepidoptera | - | -0.5539 | 1.242 | 1.662 | 0.117 |
| Lithobiomorpha | - | -0.327 | 1.083 | 2.058 | 0.121 |
| Neuroptera | - | -0.515 | 1.251 | 1.533 | 0.152 |
| Odonata | - | -0.588 | -0.386 | 4.438 | 0.181 |
| Opiliones | - | -0.243 | 1.442 | 1.262 | 0.132 |
| Orthoptera | - | -0.095 | 0.968 | 1.729 | 0.109 |
| Polydesmida | - | -0.417 | 1.245 | 1.809 | 0.141 |
| Pseudoscorpionida | - | -0.801 | 1.750 | 0.300 | 0.142 |
| Psocoptera | - | -0.936 | 2.294 | 0.666 | 0.235 |
| Scolopendromorpha | - | -0.962 | 1.669 | 0.300 | 0.051 |
| **Model 3: Length-Width-Geographic region (LWR)** |  |  |  |  |  |
| - | temperate | -0.281 | 1.030 | 1.597 | 0.149 |
| - | tropical | -0.370 | 1.086 | 1.649 | 0.186 |
| **Model 4: Length-Width (LW)** |  |  |  |  |  |
| - | - | -0.339 | 1.066 | 1.640 | 0.172 |
| **Model 5: Length-Taxonomic group-Geographic region (LTR)** |  |  |  |  |  |
| Araneae | temperate | -0.733 | 2.623 | - | 0.151 |
| Coleoptera | temperate | -0.935 | 2.455 | - | 0.295 |
| Dermaptera | temperate | -0.947 | 2.337 | - | 0.114 |
| Diptera | temperate | -1.057 | 2.489 | - | 0.182 |
| Hemiptera | temperate | -0.902 | 2.386 | - | 0.219 |
| Hymenoptera | temperate | -1.486 | 3.018 | - | 0.195 |
| Isopoda | temperate | -1.292 | 2.950 | - | 0.110 |
| Lepidoptera | temperate | -1.294 | 2.493 | - | 0.194 |
| Lithobiomorpha | temperate | -1.671 | 2.780 | - | 0.101 |
| Neuroptera | temperate | 0.156 | 0.889 | - | 0.169 |
| Opiliones | temperate | -0.364 | 2.379 | - | 0.157 |
| Orthoptera | temperate | -0.640 | 2.267 | - | 0.158 |
| Polydesmida | temperate | -1.519 | 2.595 | - | 0.037 |
| Araneae | tropical | -0.862 | 2.611 | - | 0.191 |
| Coleoptera | tropical | -1.104 | 2.553 | - | 0.375 |
| Dermaptera | tropical | -1.775 | 2.929 | - | 0.135 |
| Dictyoptera | tropical | -0.644 | 1.913 | - | 0.303 |
| Diptera | tropical | -0.973 | 2.270 | - | 0.249 |
| Geophilomorpha | tropical | -2.917 | 2.837 | - | 0.225 |
| Hemiptera | tropical | -0.813 | 2.189 | - | 0.213 |
| Hymenoptera | tropical | -1.422 | 2.792 | - | 0.248 |
| Isopoda | tropical | -1.268 | 2.839 | - | 0.124 |
| Lepidoptera | tropical | -1.433 | 2.587 | - | 0.251 |
| Lithobiomorpha | tropical | -1.884 | 2.701 | - | 0.166 |
| Neuroptera | tropical | -0.884 | 2.112 | - | 0.197 |
| Odonata | tropical | -0.816 | 1.856 | - | 0.300 |
| Opiliones | tropical | -0.453 | 2.648 | - | 0.129 |
| Orthoptera | tropical | -0.775 | 2.205 | - | 0.192 |
| Polydesmida | tropical | -1.825 | 2.726 | - | 0.184 |
| Pseudoscorpionida | tropical | -0.942 | 2.015 | - | 0.149 |
| Psocoptera | tropical | -1.154 | 2.710 | - | 0.237 |
| Scolopendromorpha | tropical | -2.084 | 2.702 | - | 0.116 |
| **Model 6: Length-Taxonomic group (LT)** |  |  |  |  |  |
| Araneae | - | -0.830 | 2.637 | - | 0.190 |
| Coleoptera | - | -1.042 | 2.537 | - | 0.334 |
| Dermaptera | - | -1.316 | 2.529 | - | 0.206 |
| Dictyoptera | - | -0.644 | 1.913 | - | 0.303 |
| Diptera | - | -1.0318 | 2.430 | - | 0.205 |
| Geophilomorpha | - | -2.917 | 2.837 | - | 0.225 |
| Hemiptera | - | -0.817 | 2.237 | - | 0.219 |
| Hymenoptera | - | -1.401 | 2.809 | - | 0.235 |
| Isopoda | - | -1.322 | 2.967 | - | 0.119 |
| Lepidoptera | - | -1.390 | 2.554 | - | 0.24 |
| Lithobiomorpha | - | -1.888 | 2.934 | - | 0.169 |
| Neuroptera | - | -0.871 | 2.010 | - | 0.217 |
| Odonata | - | -0.816 | 1.856 | - | 0.300 |
| Opiliones | - | -0.385 | 2.439 | - | 0.154 |
| Orthoptera | - | -0.791 | 2.245 | - | 0.199 |
| Polydesmida | - | -1.986 | 2.944 | - | 0.175 |
| Psocoptera | - | -1.154 | 2.710 | - | 0.237 |
| Pseudoscorpionida | - | -0.942 | 2.015 | - | 0.149 |
| Scolopendromorpha |  | -2.084 | 2.702 | - | 0.116 |
| **Model 7: Length-Geographic region (LR)** |  |  |  |  |  |
| - | temperate | -0.730 | 2.175 | - | 0.283 |
| - | tropical | -0.822 | 2.146 | - | 0.327 |
| **Model 8: Length (L)** |  |  |  |  |  |
| - | - | -0.786 | 2.166 | - | 0.313 |

Regression equations for the eight models:
Model 1 (LWTR): $\log_{10}(\text{body mass}) = a_{\text{taxon region}} + b_{\text{length taxon region}} \times \log_{10}(\text{body length}) + b_{\text{width taxon region}} \times \log_{10}(\text{body width})$
Model 2 (LWT): $\log_{10}(\text{body mass}) = a_{\text{taxon}} + b_{\text{length taxon}} \times \log_{10}(\text{body length}_{\text{taxon}}) + b_{\text{width taxon}} \times \log_{10}(\text{body width})$
Model 3 (LWR): $\log_{10}(\text{body mass}) = a_{\text{region}} + b_{\text{length region}} \times \log_{10}(\text{body length}) + b_{\text{width region}} \times \log_{10}(\text{body width})$
Model 4 (LW): $\log_{10}(\text{body mass}) = a + b_{\text{length}} \times \log_{10}(\text{body length}) + b_{\text{width}} \times \log_{10}(\text{body width})$
Model 5 (LTR): $\log_{10}(\text{body mass}) = a_{\text{taxon region}} + b_{\text{taxon region}} \times \log_{10}(\text{body length})$
Model 6 (LT): $\log_{10}(\text{body mass}) = a_{\text{taxon}} + b_{\text{taxon}} \times \log_{10}(\text{body length})$
Model 7 (LR): $\log_{10}(\text{body mass}) = a_{\text{region}} + b_{\text{region}} \times \log_{10}(\text{body length})$
Model 8 (L): $\log_{10}(\text{body mass}) = a + b \times \log_{10}(\text{body length})$

**Figure 1:**
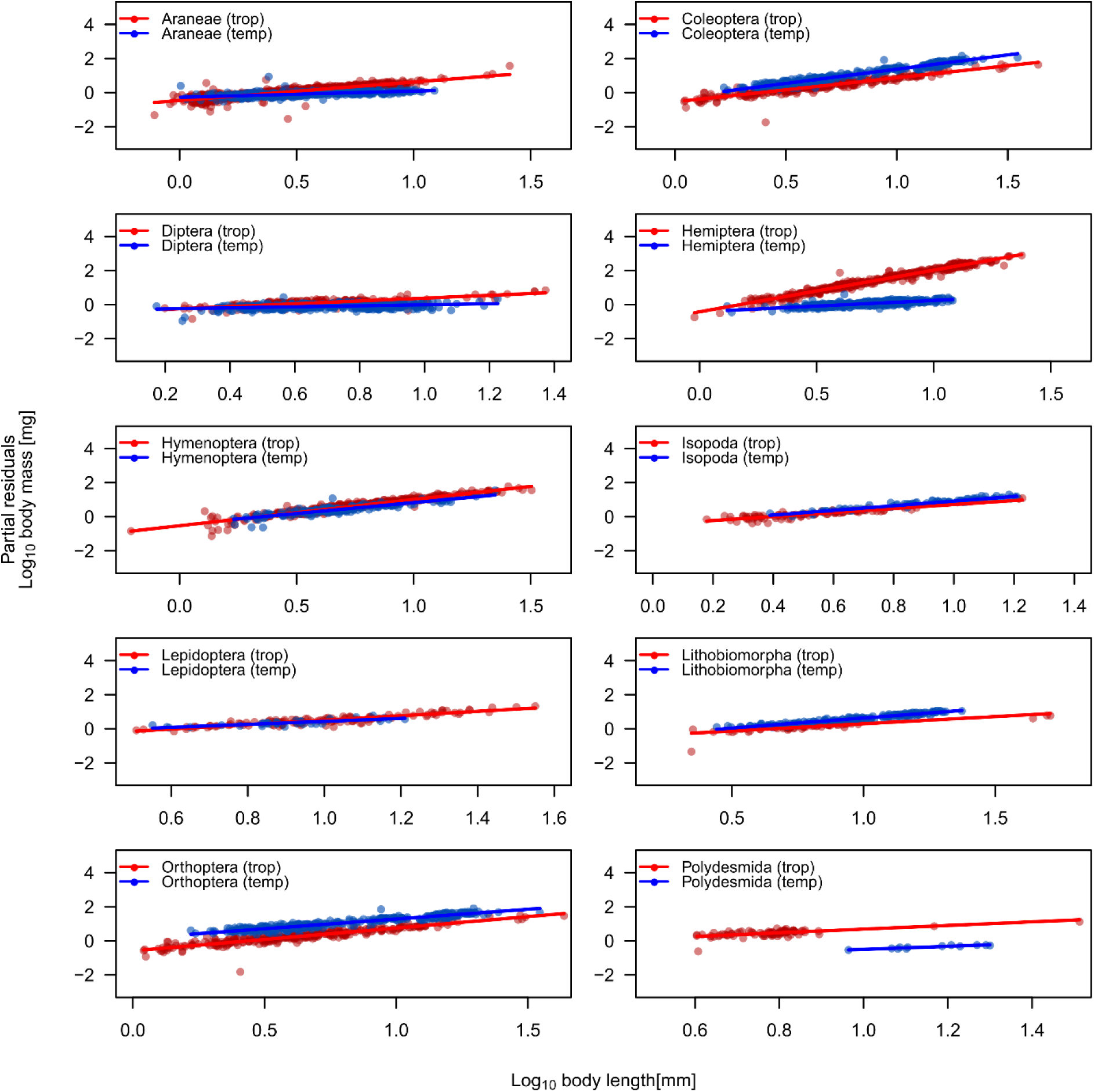
Length-mass regressions of the best fit model, which included body length, maximum body width, taxonomy and geographic region (LWTR) to predict body mass for the ten most abundant arthropod groups from the temperate (blue) and tropical (red) study areas. The y-axis displays partial residuals and, therefore, shows the effect of body length after correcting for the other variables.

The eight different models explained between 81.4 % (model L, least complex model) and 98.6 % (model LWTR, most complex model) of the total variance in body mass (Table 1). According to AIC comparisons, the four models that included body width as a co-variate explained more variation in body mass than models that only included body length as a predictor. In contrast to the results from AIC comparisons, however, r^2-^values suggested that the model including taxonomic group but not body width (model LTR and model LT, Table 1) explained marginally more variance in body mass than the model including body width but not taxonomic group or geographic region (model LW, Table 1).

Finally, to test if the application of geographically-disjunct regression parameters (i.e., where regression parameters obtained from one geographic region are used to predict body mass of arthropods in a different geographic region) increases error in body mass predictions, we calculated body mass using geographically-disjunct and geographically non-disjunct regression parameters and quantified the difference from observed body mass. In general, we found that the application of geographically-disjunct parameters for whole-fauna regressions led to increased prediction error of body mass when compared to using non-disjunct regression parameters (Figure 2). Whether this prediction error leads to an under- or over-estimation of body mass depended on the geographic region and the morphological traits used to predict body mass. With only body length included as a predictor (Model LR), body mass of temperate arthropods was underestimated on average by 33 % (geometric-mean ratio = 0.77) using tropical regression parameters (Figure 2a), whereas tropical arthropod body mass was overestimated on average by 29 % (geometric-mean ratio = 1.29) when using temperate regression parameters (Figure 2b). Interestingly, when using model LR, prediction error increased with increasing body length for both temperate and tropical arthropods using geographically-disjunct regression parameters (Figure 2a, b). In contrast, when body width was included in the model, the geographically-disjunct regression prediction error shifted between overestimation and underestimation with increasing body length. For temperate arthropods, the models tended to underestimate predicted body mass at small body lengths and overestimate predicted body mass at large body lengths, with an average underestimation of 8% (geometric-mean ratio = 0.92) (Figure 2c). In contrast, body mass of tropical arthropods was overestimated at smaller body lengths and underestimated at larger body lengths when using geographically-disjunct regression parameters in model LWR, with an average overestimation of 10 % (geometric-mean ratio = 1.10) (Figure 2d).

**Figure 2:**
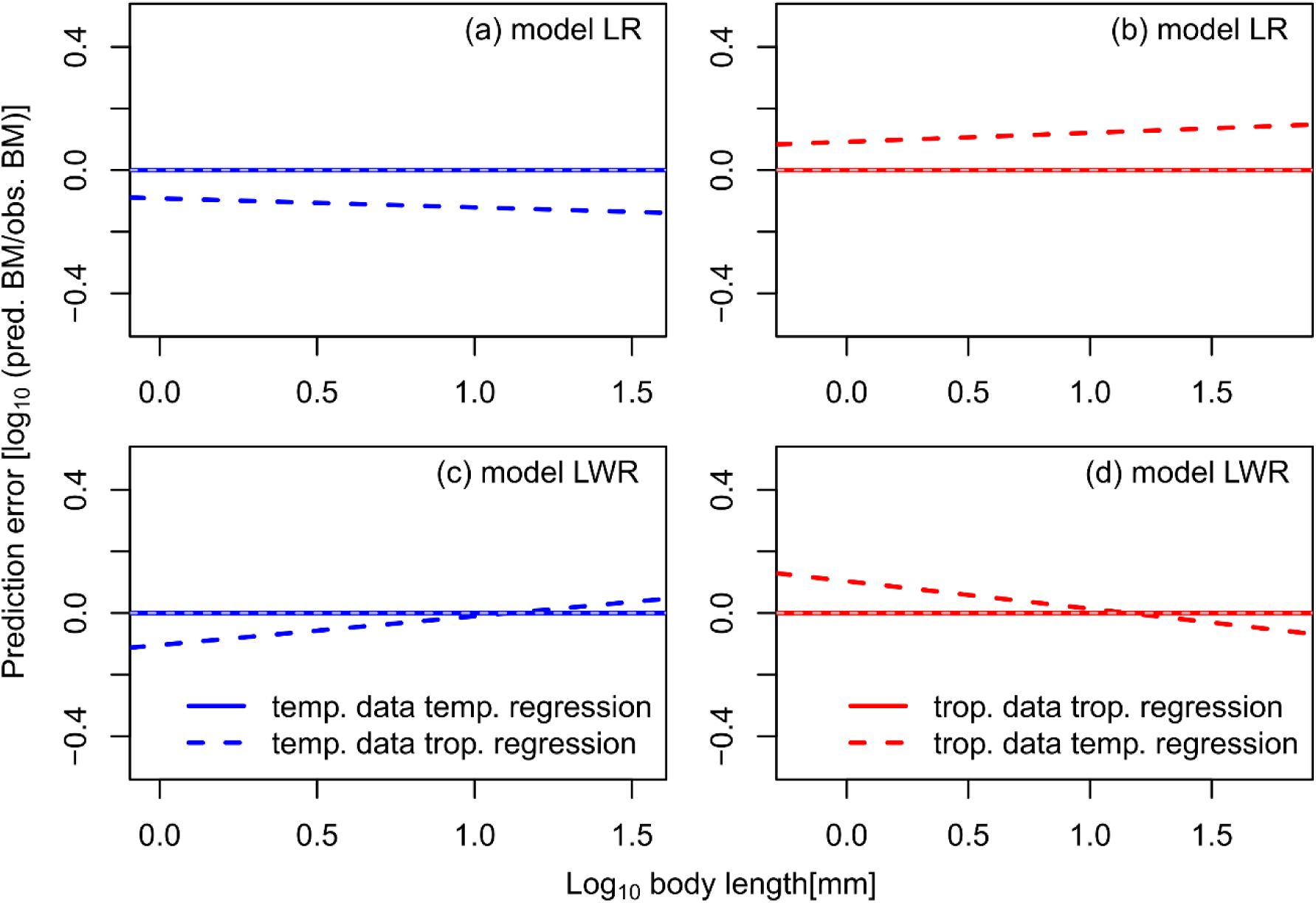
Prediction error (log response ratio of predicted versus observed body mass values) for temperate (blue lines, panels a and c) and tropical (red lines, panel’s b and d) arthropod body mass obtained by using geographically disjunct (dashed lines) and non-disjunct (solid lines) regression parameters for the LR (a and b) and LWR (c and d) models. LR = length + region and LWR = length + width + region models.

## Discussion

A wide range of individual-to community-level characteristics are influenced by body size, including abundance, metabolic rate, movement speed or growth rate (Gillooly *et al.*, 2001; White *et al.*, 2007; Hirt *et al.*, 2017). In order to make realistic predictions of these measures, it is essential to have reliable body mass data of target organisms. In our dataset consisting of 6293 organisms spanning 19 higher order taxa from both tropical and temperate geographic regions, we found an overall positive power law relationship between body mass and body length across taxonomic groups and the tropical and temperate geographic regions. The only exception to this universal trend was for Odonata and Neuroptera, which showed a negative relationship between body mass and body length in a subset of models.

The slope of the relationship between body length and mass depended on taxonomic group and geographic region of arthropods. Furthermore, adding body width as an additional morphological predictor strongly improved body mass prediction accuracy. This is probably due to certain groups where the body length-to-width ratio is considerably different to the average of all taxonomic groups (e.g., Staphyilinid beetles have a higher body length-to-width ratio than other beetle families). Thus, using body length as the only predictor of body mass is almost certainly insufficient to capture the morphological variation present within taxonomic groups. Therefore, we expected that the incorporation of body width as an additional predictor in our models should increase the accuracy of body mass predictions. Consistent with our expectations, we found that including body width into the estimation of body mass resulted in a strong improvement of prediction accuracy, in comparison to using body length, alone, as a single predictor of body mass. Moreover, incorporating only body width as an additional predictor yielded higher prediction accuracy than incorporating taxonomic group and geographic region into the models. Body mass is related to the volume of an organism, which can be described by length, width and height. Hence, adding height to predict body mass could lead to more accurate body mass estimations than using only body length and width. Measuring another morphological trait of an organism, however, increases time needed for processing samples, presenting a trade-off between maximising prediction accuracy and minimising time spent measuring traits. As more than 98 % of variance in body mass was described by length, width, taxonomic group and geographic region, the benefit of adding body height would unlikely outweigh the added workload. Indeed, previous studies have shown that including body shape (i.e. body length and width) instead of taxonomy lead to more accurate body mass estimates at the order level, but not at higher taxonomic resolution (Gruner, 2003; Wardhaugh, 2013). Our results strongly support the finding that the accuracy in predicting body mass improves with additional morphological traits in addition to body length for scaling relationships conducted at the order level.

In addition to body width, taxonomic group and geographic origin of the arthropods also influenced the relationship between body length and body mass. This is likely because variation in arthropod body size is influenced by a range of other factors such as evolutionary history and environmental variation (Chown & Gaston, 2010). For example, Bergmann’s rule proposes that body size increases with latitude, though the opposite has been observed for arthropods (Mousseau, 1997). Generally speaking, these concepts suggest that the body size of arthropods depends strongly on their geographic origin, particularly with respect to latitude. Therefore, we expected that the application of geographically-disjunct regression parameters from tropical and temperate regions could lead to significant prediction error in arthropod body mass. If researchers are unable to use regression parameters from data collected in a similar geographic regions to their study site (due to a lack of available scaling relationships), this could have important consequences for the body mass-related results drawn from such studies. Consistent with our expectations, we found that the use of geographically-disjunct length-mass regression parameters led to inaccurate body mass predictions ranging between average prediction-errors of 8 % to 33 %, depending on the model used. Furthermore, when only body length was used as a morphological predictor, body mass prediction accuracy of geographically-disjunct regressions decreased with increasing body length of arthropods. This has important consequences for the quality of body mass data, as our results suggest that body mass of longer arthropods will be more severely over- or underestimated than that of shorter arthropods. Therefore, our results highlight a potential systematic bias of decreasing prediction accuracy with increasing body length when applying regression parameters from different geographical regions. Ultimately, studies investigating body size responses to environmental conditions and the resulting impacts on ecosystem functioning rely on accurate calculations of body mass. Therefore, it is essential for such studies to use length-mass regression parameters that are obtained from similar geographic origins as the organisms for which body mass is being predicted.

Our study provides a highly comprehensive set of regression parameters for predicting live body mass of terrestrial arthropods. This set of regression parameters is useful for researchers wishing to quantify body mass of arthropods across a range of underlying morphological traits, taxonomic identities, and geographical regions. By incorporating all combinations of geographic region, taxonomic group and body width in our allometric models, our results allow investigators to choose length-mass regression parameters for predicting body mass across a broad variety of arthropod datasets. Additionally, we provide an explicit estimation of the prediction error caused by using geographically disjunct regression parameters, to assist in deciding which regression parameters will be the most appropriate for predicting arthropod body mass for a given dataset. In summary, our results will aid future studies in accurately assessing body mass of arthropods, thus increasing our ability to further explore the ecological implications of body size.

## Acknowledgements

This study was financed by the Deutsche Forschungsgemeinschaft (DFG) in the framework of the collaborative German-Indonesian research project CRC990, phase I. Tropical invertebrates were collected based on Permit No. 2695/IPH.1/KS.02/XI/2012 recommended by the Indonesian Institute of Sciences (LIPI) and issued by the Ministry of Forestry (PHKA). ADB, MJ, and UB were supported by the German Research Foundation within the framework of the Jena Experiment (FOR 1451). ES was supported by the German Federal Environmental Foundation (DBU). Further, MJ was supported by the Swiss National Science Foundation. We thank Nisfi Yuniar, Windi Ayu Prawitasari, Paul Brugman, Theresa Bodner, Franca Marian and Kristina Barnes for their help with sampling, sorting and identification of arthropods in Indonesia and Germany.

## References

Barnes, A. D., Jochum, M., Lefcheck, J. S., Eisenhauer, N., Scherber, C., O’Connor, M. I., de Ruiter, P. & Brose, U. (2018) Energy Flux: The Link between Multitrophic Biodiversity and Ecosystem Functioning. Trends in Ecology & Evolution, 33, 186–197.

Bekoff, M., Diamond, J. & Mitton, J. B. (1981) Life-history patterns and sociality in canids: Body size, reproduction, and behavior. Oecologia, 50, 386–390.

Belgrano, A., Allen, A. P., Enquist, B. J. & Gillooly, J. F. (2002) Allometric scaling of maximum population density: a common rule for marine phytoplankton and terrestrial plants. Ecology Letters, 5, 611–613.

Benke, A. C., Huryn, A. D., Smock, L. A. & Wallace, J. B. (1999) Length-Mass Relationships for Freshwater Macroinvertebrates in North America with Particular Reference to the Southeastern United States. Journal of the North American Benthological Society, 18, 308–343.

Boudreau, P. R., Dickie, L. M. & Kerr, S. R. (1991) Body-size spectra of production and biomass as system-level indicators of ecological dynamics. Journal of Theoretical Biology, 152, 329–339.

Brose, U., Williams, R. J. & Martinez, N. D. (2006) Allometric scaling enhances stability in complex food webs. Ecology letters, 9, 1228–36.

Brown, J. H., Gillooly, J. F., Allen, A. P., Savage, V. M. & West, G. B. (2004) Toward a metabolic theory of ecology. Ecology, 85, 1771–1789.

Burgherr, M. & Meyer, E. I. (1997) Regression analysis of linear body dimensions vs. dry mass in stream macroinvertebrates. Archiv für Hydrobiologie, 139, 101–112.

Chown, S. L. & Gaston, K. J. (2010) Body size variation in insects: a macroecological perspective. Biological Reviews, 85, 139–169.

Commonwealth Scientific and Industrial Research Organization (Australia) (1991) The insects of Australia: A Textbook for Students and Research Workers (2 Volume Set). Cornell University Press, Ithaca, USA.

Ehnes, R. B., Rall, B. C. & Brose, U. (2011) Phylogenetic grouping, curvature and metabolic scaling in terrestrial invertebrates. Ecology Letters, 14, 993–1000.

Gillooly, J. F., Brown, J. H., West, G. B., Savage, V. M. & Charnov, E. L. (2001) Effects of Size and Temperature on Metabolic Rate. Science, 293.

Greenleaf, S. S., Williams, N. M., Winfree, R. & Kremen, C. (2007) Bee foraging ranges and their relationship to body size. Oecologia, 153, 589–596.

Gruner, D. S. (2003) Regressions of length and width to predict arthropod biomass in the Hawaiian Islands. Pacific Science, 57, 325–336.

Heinrichs, S., Winterhoff, W. & Schmidt, W. (2014) 50 years of constancy and dynamics of calcareous beech forests on dry slopes (Carici-Fagetum) - A comparison of old and recent vegetation releves from the Gottingen forest. TUEXENIA, 34, 9–38.

Hirt, M. R., Jetz, W., Rall, B. C. & Brose, U. (2017) A general scaling law reveals why the largest animals are not the fastest. Nature Ecology & Evolution, 1, 1116–1122.

Ishizuka, S., Tsuruta, H. & Murdiyarso, D. (2002) An intensive field study on CO2, CH4, and N2O emissions from soils at four land-use types in Sumatra, Indonesia. Global Biogeochemical Cycles, 16, 22-1–22–11.

Jetz, W., Carbone, C., Fulford, J. & Brown, J. H. (2004) The Scaling of Animal Space Use. Science, 306, 266–268.

Jocqué, R. & Dippenaar-Schoeman, A. (2007) Spider families of the world. Royal Museum for Central Africa, Tervuren, Belgium.

Johnston, T. A. & Cunjak, R. A. (1999) Dry mass-length relationships for benthic insects: a review with new data from Catamaran Brook, New Brunswick, Canada. Freshwater Biology, 41, 653–674.

Kalinkat, G., Brose, U. & Rall, B. C. (2013) Habitat structure alters top-down control in litter communities. Oecologia, 172, 877–887.

Kleiber, M. (1932) Body size and metabolism. Hilgardia, 6, 315–351.

Kolokotrones, T., Van Savage, Deeds, E. J. & Fontana, W. (2010) Curvature in metabolic scaling. Nature, 464, 753–756.

Lindstedt, S. L., Miller, B. J. & Buskirk, S. W. (1986) Home range, time, and body size in mammals. Ecology. Ecological Society of America, 67, 413–418.

Mercer, R. D., Gabriel, A. G. A., Barendse, J., Marshall, D. J. & Chown, S. L. (2001) Invertebrate body sizes from Marion Island. Antarctic Science, 13, 135–143.

Mousseau, T. A. (1997) Ectotherms Follow the Converse to Bergmann’s Rule. Evolution, 51, 630–632.

Peters, R. H. (1983) The ecological implications of body size. Cambridge University Press, Cambridge, UK.

R Core Team (2015) R: a language and environment for statistical computing. Vienna, Austria: R Foundation for Statistical Computing.

Rall, B. C., Brose, U., Hartvig, M., Kalinkat, G., Schwarzmuller, F., Vucic-Pestic, O. & Petchey, O. L. (2012) Universal temperature and body-mass scaling of feeding rates. Philosophical Transactions of the Royal Society B: Biological Sciences, 367, 2923–2934.

Reichle, D. (1968) Relation of Body Size to Food Intake, Oxygen Consumption, and Trace Element Metabolism in Forest Floor Arthropods. Ecology. Ecological Society of America, 49, 538–542.

Riede, J. O., Brose, U., Ebenman, B., Jacob, U., Thompson, R., Townsend, C. R. & Jonsson, T. (2011) Stepping in Elton’s footprints: a general scaling model for body masses and trophic levels across ecosystems. Ecology Letters, 14, 169–178.

Rogers, L. E., Buschbom, R. L. & Watson, C. R. (1977) Length-Weight Relationships of Shrub-Steppe Invertebrates. Annals of the Entomological Society of America, 70, 51–53.

Schaefer, M. (2009) Brohmer - Fauna von Deutschland. Quelle & Meyer, Wiebelsheim, Germany.

Schoener, T. W. (1980) Length-Weight Regressions in Tropical and Temperate Forest-Understory Insects. Annals of the Entomological Society of America, 73, 106–109.

Swihart, R. K., Slade, N. A. & Bergstrom, B. J. (1988) Relating Body Size to the Rate of Home Range Use in Mammals. Ecology. Ecological Society of America, 69, 393–399.

Violle, C., Enquist, B. J., McGill, B. J., Jiang, L., Albert, C. H., Hulshof, C., Jung, V. & Messier, J. (2012) The return of the variance: intraspecific variability in community ecology. Trends in Ecology & Evolution, 27, 244–252.

Wardhaugh, C. W. (2013) Estimation of biomass from body length and width for tropical rainforest canopy invertebrates. Australian Journal of Entomology, 52, 291–298.

White, E. P., Ernest, S. K. M., Kerkhoff, A. J. & Enquist, B. J. (2007) Relationships between body size and abundance in ecology. Trends in Ecology & Evolution, 22, 323–330.

Woodward, G., Ebenman, B., Emmerson, M., Montoya, J., Olesen, J., Valido, A. & Warren, P. (2005) Body size in ecological networks. Trends in Ecology & Evolution, 20, 402–409.

